# A practical workflow for CAR screening in primary T cells using fitness-guided design and mRNA electroporation

**DOI:** 10.64898/2026.07.28.741207

**Authors:** Atsushi Okuma, Yoshihito Ishida, Shoji Hisada

## Abstract

Chimeric antigen receptor (CAR) T cell therapy has achieved remarkable therapeutic outcomes in hematological cancers. However, broader clinical use has uncovered substantial challenges arising from intrinsic properties of both T cells and tumor tissues. As the functional phenotype of CAR T cells is affected by the CAR molecular architecture, optimizing CAR constructs continues to be a critical and ongoing task. Here, we present a practical workflow for scalable screening of CAR variants in primary T cells using fitness-guided design and mRNA electroporation. Using a CD19-targeted second-generation CAR, we built a library of point mutants that focused mutagenesis on hinge and costimulatory domains. Amino acid substitutions were prioritized using the sequence-based zero-shot fitness predictor to enrich evolutionarily tolerated variants. From 340 designed variants, we electroporated mRNA encoding 85 constructs into primary human CD8+ T cells and quantified cytotoxicity against CD19-positive Nalm6 cells. Twenty-four variants reproducibly exceeded wild-type cytotoxicity across three runs, and three hits were selected for lentiviral validation. One of the selected variants showed significantly improved cytotoxicity despite lower expression frequency and exhibited higher CD62L within CAR-positive cells, suggesting enhanced intrinsic function with a less differentiated phenotype. This approach enables scalable, rapid discovery of improved CAR domain variants directly in primary T cells.

## Introduction

Chimeric antigen receptor (CAR) T cell therapy has transformed the treatment of B-cell leukemia and lymphoma. CARs are synthetic receptors typically composed of five components (Roex et al., 2025): (1) an antigen-binding domain (ABD), (2) a hinge domain (HD), (3) a transmembrane domain (TMD), (4) one or more costimulatory domains (CSDs), and (5) a T cell activation domain (TAD). While ABDs have been extensively diversified and engineered—spanning dozens of target antigens (MacKay et al., 2020) and approaches such as affinity tuning (Drent et al., 2017) and alternative binding scaffolds (Ellebrecht et al., 2016; Lam et al., 2020; Lee et al., 2023; McComb et al., 2022; Perriello et al., 2023; Sauer et al., 2021; Stepanov et al., 2018; Van Den Eynde et al., 2024)—the remaining CAR modules used clinically are comparatively constrained. Most approved CARs rely on a limited set of domains (e.g., CD8α-, CD28-, or IgG4-derived HDs; CD8α- or CD28-derived TMDs; CD28- or 4-1BB-derived CSDs; and CD3ζ-derived TAD) (Mitra et al., 2023).

To efficiently explore CAR design space, high-throughput CAR screening approaches have recently emerged (Alonso-Camino et al., 2013; Bloemberg et al., 2020; Butler et al., 2023; Castellanos-Rueda et al., 2022; Daniels et al., 2022; Duong et al., 2013; Fierle et al., 2022; Fu et al., 2022; Goodman et al., 2022; Gordon et al., 2022; Lipowska-Bhalla et al., 2013; Ma et al., 2021; Ochi et al., 2021; Rios et al., 2023; Roberto et al., 2020; Rydzek et al., 2019). Many of these approaches have utilized pooled libraries to maximize scale, coupling genotype to phenotype through enrichment or selection. However, pooled library formats impose stringent requirements on gene delivery and library representation in T cells, and phenotypes can be difficult to attribute to individual constructs when multiple variants enter the same cell.

A central technical challenge in CAR library screening is gene delivery. The pooled library approaches generally require stable, single-construct integration per cell to avoid co-selection artifacts. Therefore, viral transduction (Alonso-Camino et al., 2013; Duong et al., 2013; Elazar et al., 2022; Fierle et al., 2022; Fu et al., 2022; Goodman et al., 2022; Gordon et al., 2022; Krokhotin et al., 2019; Lipowska-Bhalla et al., 2013; Ma et al., 2021; Ochi et al., 2021; Rios et al., 2023), transposon systems (Rydzek et al., 2019), or targeted genome editing (Castellanos-Rueda et al., 2022; Roberto et al., 2020) are commonly used. In contrast, transient delivery methods such as electroporation of CAR-encoding nucleic acids are often considered less suitable for pooled enrichment screens because high intracellular concentrations of mixed library elements can result in multiple constructs per cell and confound selection (Lipowska-Bhalla et al., 2013). Nevertheless, for arrayed screening—where each construct is evaluated independently rather than enriched from a pool—transient expression can provide a rapid and scalable alternative to viral transduction, enabling parallel functional testing of many CAR designs directly in primary T cells. In this context, mRNA electroporation offers a rapid, virus-free method to generate CAR T cells, enabling short design–build–test cycles and side-by-side comparison of many constructs under matched donor and culture conditions.

Here, we developed an arrayed screening strategy to identify improved CAR variants by focusing mutagenesis on hinge and costimulatory domains while preserving core receptor functions. Using a CD19-directed second-generation CAR as a template, we constructed a library of CAR variants incorporating amino acid substitutions prioritized by predicted protein fitness using a sequence-based zero-shot model. We then implemented mRNA electroporation into primary CD8+ T cells to rapidly screen cytotoxic activity against CD19-positive Nalm6 cells and validated the screening hits using lentiviral transduction. This work establishes a practical workflow for scalable, primary T cell–based screening of CAR domain variants.

## Results

### Design of a CAR variant library

To establish a practical strategy for expanding CAR sequence diversity, we designed CAR variants incorporating amino acid substitutions. We selected a second-generation CAR as the template, consisting of an FMC63 scFv antigen-binding domain, CD28-derived hinge, transmembrane, and costimulatory domains, and a CD3ζ T cell activation domain (hereafter 28z CAR). To minimize the generation of non-functional CAR molecules in T cells, we preserved domains required for antigen binding, membrane localization, and minimal intracellular signaling, and therefore confined mutagenesis to the hinge and costimulatory domains. In this design, we introduced at most one amino acid substitution per domain (i.e., one substitution in the hinge and/or one substitution in the costimulatory domain) to reduce the likelihood of generating non-functional constructs.

To reduce the frequency of variants with no or extremely low function, we prioritized amino acid substitutions with higher protein fitness predicted by EVmutation (Hopf et al., 2017). EVmutation is a sequence-based zero-shot predictor that relies on local sequence information. It searches sequence databases to build a multiple sequence alignment (MSA) for a target sequence and then learns a representation of the underlying sequence distribution that constrains local protein sequences. To predict fitness across as many positions as possible, we prioritized MSAs with higher positional coverage rather than maximizing the number of sequences. We generated MSAs containing 2,480 and 73 redundancy-reduced sequences that covered 82% (32/39) and 90% (37/41) of positions in the hinge and costimulatory domain, respectively. For the covered regions, we obtained single-mutant matrices and ranked substitutions by their evolutionary statistical energy relative to the corresponding wild-type amino acid (ΔE), interpreting higher ΔE as favorable (higher predicted fitness). Notably, substitutions introduced into the CD28 costimulatory domain were selected from positions outside canonical signaling motifs (YMNM, PRRP, and PYAP), thereby avoiding direct perturbation of these well-characterized interaction sites. Including the wild-type sequence for each domain, we designed 17 hinge domain sequences and 20 costimulatory domain sequences (Fig. 1). By combining these sequences, we generated a total of 340 CAR variants.

**Fig. 1.**
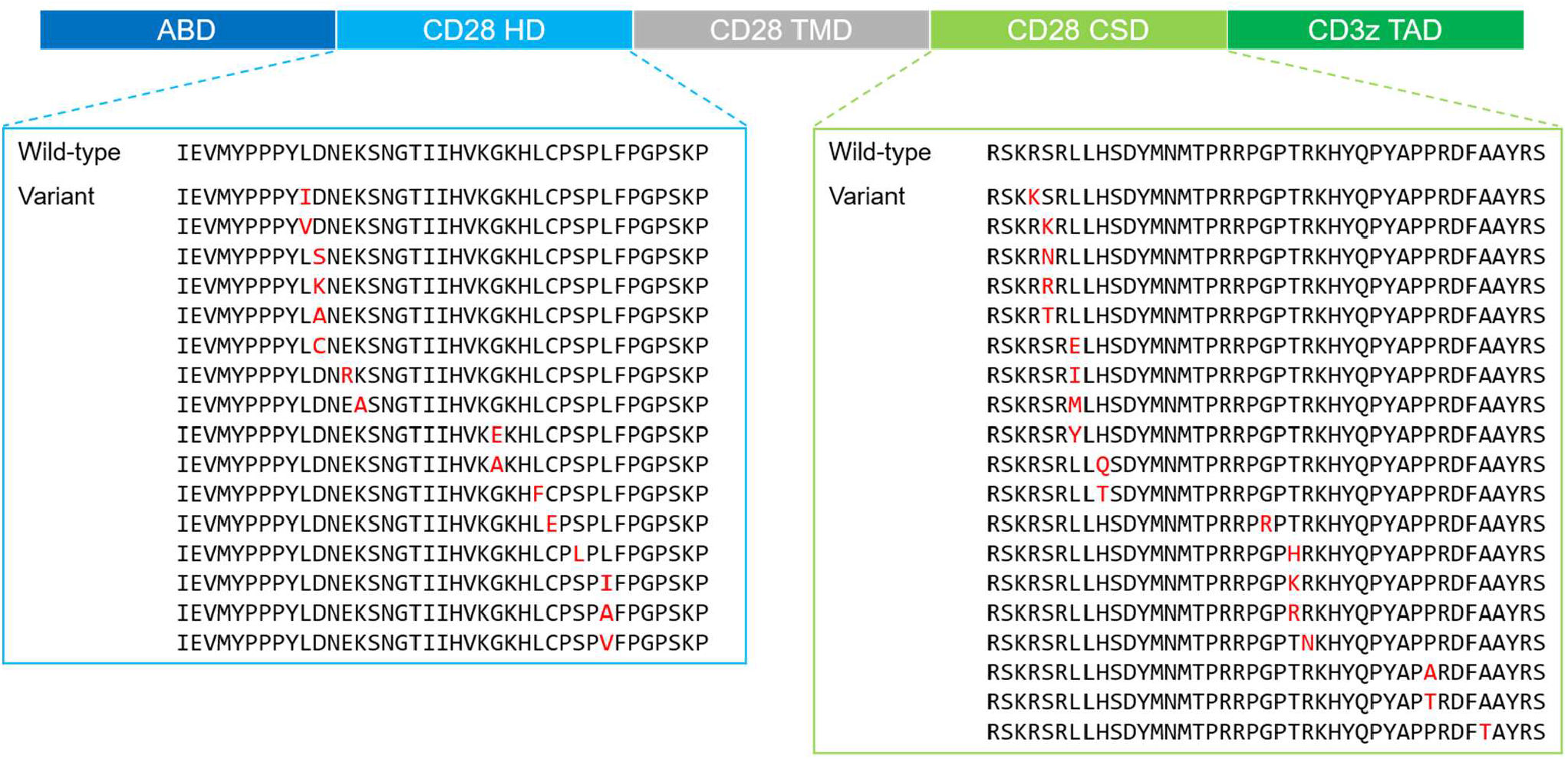
Schematic representation of a CAR containing an antigen-binding domain (ABD), a transmembrane domain (TMD), a costimulatory domain (CSD), and a T cell activation domain (TAD). The wild-type and variant sequences of the CD28-derived hinge and costimulatory domains used to build the combinatorial library are shown. Each variant sequence contains single amino acid substitution (highlighted in red) prioritized by predicted protein fitness. In total, 17 hinge sequences and 20 costimulatory sequences (including wild-type) were designed, yielding 340 CAR variants by combinatorial assembly.

### Optimization of a miniaturized 384-well cytotoxicity assay for arrayed screening of electroporated CAR T cells

To evaluate CAR variants individually, electroporation was utilized as the gene-delivery method for CAR T cell preparation. Although viral vectors are commonly used to produce CAR T cells, they are not well suited for introducing many different CAR constructs into T cells in parallel. First, we miniaturized the cytotoxicity assay from a 96-well to a 384-well format. The cytotoxicity of CAR T cells expressing the template CAR was assessed in both formats. The virally transduced CD8+ T cells were co-cultured with luciferase-expressing Nalm6 (Nalm-Luc) cells seeded at 50,000 cells/well in 96-well plates and 10,000 cells/well in 384-well plates. After 24 hours of co-culture, residual viable Nalm-Luc cells were quantified by a luciferase assay. The 384-well assay demonstrated excellent performance, with a Z′ factor over 0.9 and a signal-to-background (S/B) ratio of 61.6, and effector-to-target (E:T) ratio–dependent luminescence (Fig. 2).

**Fig. 2.**
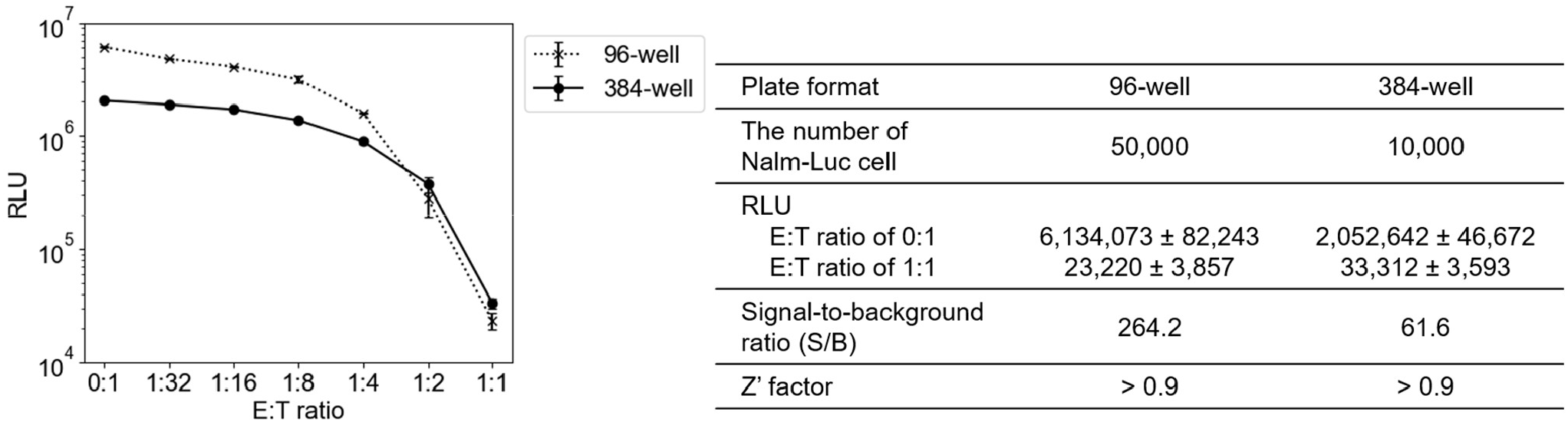
Comparison of Nalm-Luc cell luminescence in 96- and 384-well plates after 24 hours of co-culture with CD19 CAR T cells. Data are presented as mean relative light units (RLUs) ± standard deviation. Signal-to-background ratios and Z′ factors were calculated using RLUs at E:T ratios of 0:1 (no CAR T cells) and 1:1.

Next, the cytotoxicity of electroporated CAR T cells was assessed using first- and second-generation CD19 CARs with previously reported activity against Nalm6 cells (Imai et al., 2004). Electroporated T cells expressing the second-generation CAR (BBz) showed greater cytotoxicity than those expressing the first-generation CAR (z), consistent with the prior reports (Fig. 3A). For BBz, virally transduced T cells (vBBz-T) showed higher cytotoxicity than electroporated T cells (eBBz-T). Because cytotoxicity can be influenced by the number of CAR-positive cells, we used GFP co-expressed via a P2A peptide to estimate the fraction of CAR-positive cells. The frequency of GFP-positive cells was ∼45% in eBBz-T and ez-T, whereas it was ∼80% in vBBz-T (Fig. 3B). These results suggest that the enhanced cytotoxicity observed in vBBz-T is at least attributed to a higher proportion of CAR-expressing cells. Collectively, these results gave us confidence in moving forward with arrayed CAR screening using electroporation for gene delivery.

**Fig. 3.**
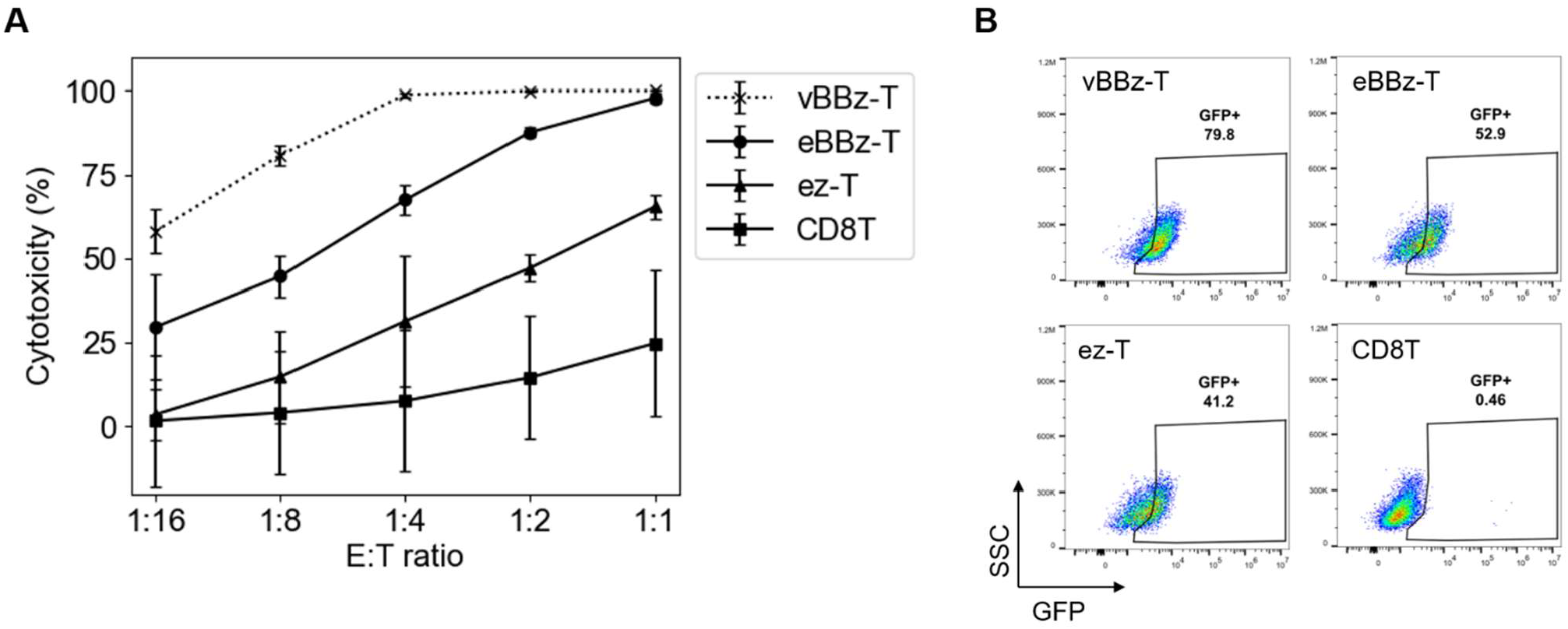
Comparison of cytotoxicity between first- and second-generation CD19 CAR T cells. Primary human CD8+ T cells (CD8T) were electroporated with mRNA encoding a first-genaration CAR (z) or a second-generation CAR (BBz) and the resultant cells (ez-T and eBBz-T) were co-cultured with Nalm-Luc cells. (A) Cytotoxicity of CD19 CAR T cells measured by a luciferase assay. vBBz-T denotes CAR T cells transduced with a lentiviral vector encoding BBz. (B) CD19 CAR expression assessed by flow cytometry. The x-axis indicates GFP co-expressed with the CAR via P2A, and the y-axis indicates side scatter (SSC).

### Identification of enhanced CAR variants

From the 340 designed CAR variants, mRNAs encoding 85 variants were introduced into CD8+ T cells by electroporation to generate CAR T cells, and cytotoxicity against Nalm-Luc cells was measured. All screens were performed at a single E:T ratio (1:2) and repeated three times. Among the 85 variants tested, 24 variants exhibited higher cytotoxicity than the wild-type CAR in all three runs. From these candidates, we selected variant #165, #306, and #327 for further evaluation (Fig. 4).

**Fig. 4.**
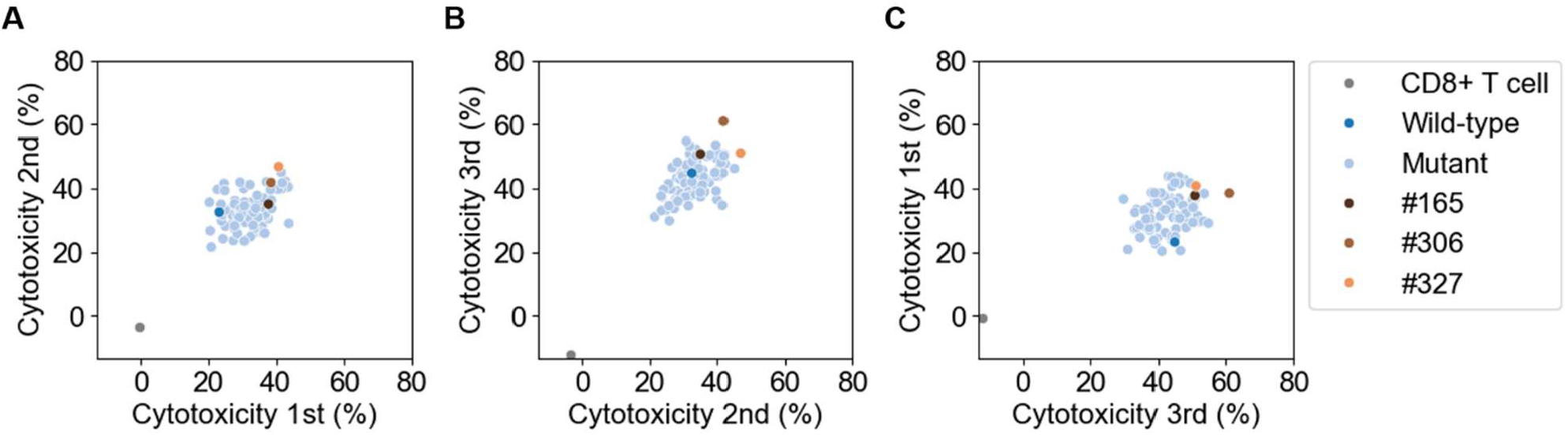
Cytotoxicity of human primary CD8+ T cells expressing wild-type and variant CAR following co-culture with Nalm-Luc cells at an effector-to-target ratio of 1:2. Remaining Nalm-Luc cells were quantified by measuring bioluminescent activity. Each dot represents one CAR construct or mock CD8+ T cells, plotted as percent killing measured in two screening runs. (A) 1st run versus 2nd run. (B) 2nd run versus 3rd run. (C) 3rd run and 1st run. Mutants exhibiting higher cytotoxicity than the wild-type CAR in all three runs (n = 24) were identified, and variants #165, #306, and #327 were selected for further evaluation.

We next generated lentivirally transduced CAR T cells expressing these three variants and the wild-type 28z CAR and compared their cytotoxicity and differentiation status (Fig. 5A, B). Because measured cytotoxicity can be affected by the frequency of CAR-expressing cells, we enriched transduced cells by sorting for GFP, which was linked to the CAR via a P2A peptide. GFP-positive cells were then co-cultured with Nalm-Luc cells to assess cytotoxicity. Variant #306 showed a modest but statistically significant increase in cytotoxicity relative to the wild-type, whereas #327 showed a marked increase. In contrast, #165 exhibited cytotoxicity comparable to that of the wild-type. To evaluate differentiation status, we measured CD62L expression in the GFP-positive population. CD62L mean fluorescence intensity (MFI) was significantly higher in #306 CAR T cells than in the wild-type CAR T cells. Taken together, these results suggest that #306 may represent a CAR variant with both a less differentiated phenotype and an enhanced cytotoxicity compared with the wild-type, whereas the apparently superior cytotoxicity of #327 may be associated with a more differentiated state.

**Fig. 5.**
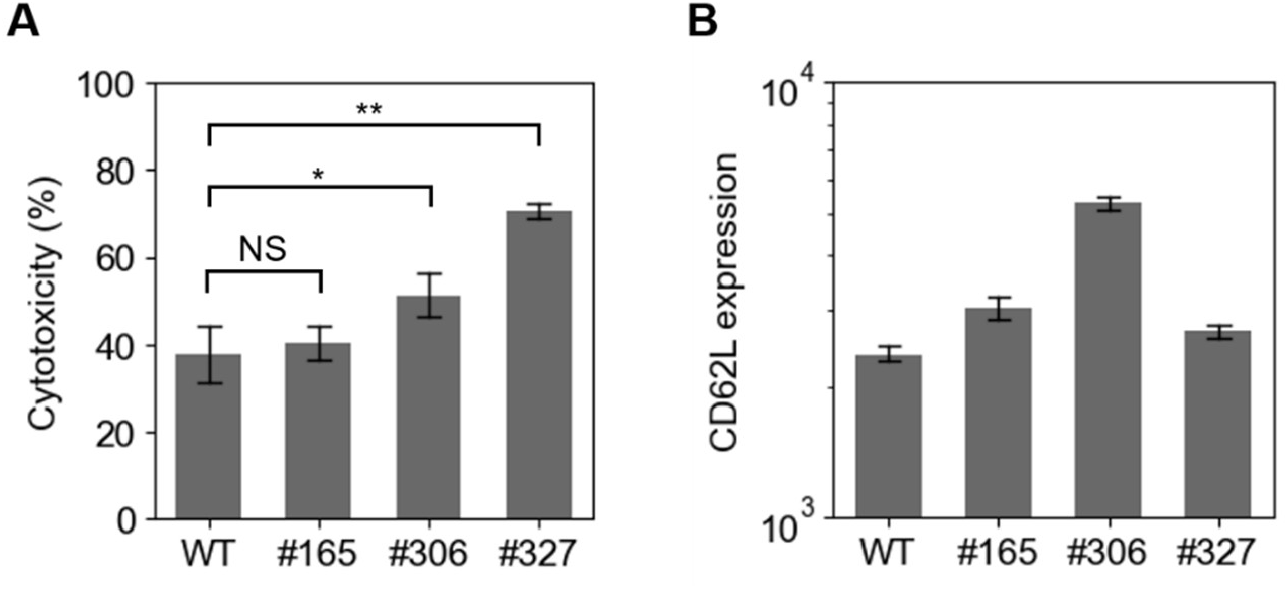
Functional and phenotypic characterization of lentivirally transduced CAR T cells expressing selected variants. Primary human CD8+ T cells were transduced with lentiviral vectors encoding the wild-type (WT) CAR or mutants #165, #306, and #327. (A) Nalm-Luc cells were co-cultured with CAR T cells at an effector-to-target ratio of 1:16, and the percentage of Nalm-Luc cell lysis were measured by bioluminescence. Data represent mean ± SD of 4 replicates from 1 independent experiment. Statistical significance was assessed using Dunnett’s multiple comparison test. * *P* < 0.001; ** *P* < 0.0001. (B) CD62L expression quantified as mean fluorescence intensity (MFI) in sorted GFP-positive T cells.

Collectively, these results indicate that screening T cells transiently expressing amino-acid–substituted CAR variants via mRNA electroporation is a feasible approach to identify CAR designs with improved cytotoxic function.

## Discussion

In this study, we developed an arrayed screening workflow to explore the mutational landscape of non-antigen-binding CAR modules and identify variants with improved functions. By restricting mutagenesis to the CD28-derived hinge and costimulatory domains while preserving the FMC63 scFv, CD28 TMD, and CD3ζ TAD, we sought to minimize the risk of generating non-functional receptors. In addition, we used EVmutation to prioritize substitutions with higher predicted protein fitness, further reducing the likelihood of severely deleterious variants, enabling efficient functional screening in primary T cells.

A key practical advantage of our approach is the use of mRNA electroporation to enable parallel testing of many CAR constructs. Viral gene transfer is the standard for CAR T cell manufacturing, but it is not well suited for rapid screening of dozens of constructs because vector production and transduction are time- and labor-intensive. In contrast, electroporation of in vitro–transcribed mRNA allowed rapid generation of CAR T cells for direct functional assessment. Importantly, we performed screening in primary CD8+ T cells rather than a reporter cell line, enabling selection based on a clinically relevant effector function (tumor cell killing). While pooled-library enrichment strategies often require stable single-copy integration to avoid multi-construct artifacts, our arrayed format evaluates each construct independently and is therefore compatible with transient expression.

Among the 85 variants tested, 24 showed higher cytotoxicity than the wild-type across all three screening runs, indicating that even single amino acid substitutions in the hinge and/or costimulatory domains can measurably tune CAR function. Notably, substitutions in the CD28 costimulatory domain were selected outside canonical signaling motifs (YMNM, PRRP, PYAP), suggesting that residues beyond well-characterized interaction sites can still modulate CAR output. These findings support the broader premise that current CAR designs—often assembled from unmodified wild-type domains—leave substantial functional sequence space unexplored.

Moreover, #306 CAR T cells displayed significantly higher CD62L MFI within the sorted GFP-positive population, indicating a less differentiated phenotype. Since less differentiated T cell states are often associated with improved proliferative potential and persistence, the combination of enhanced cytotoxicity and CD62L^high^ phenotype makes #306 a particularly interesting lead for further testing. While CD62L alone does not fully define differentiation state, the observed shift supports the possibility that #306 alters signaling strength or kinetics in a manner that preserves a more stem-like/central-memory– like profile while maintaining or improving effector function.

Several limitations should be noted. First, both the electroporation screen and lentiviral validation were performed using CD8+ T cells from a single donor, and donor-to-donor variability could affect absolute performance and variant ranking. Second, cytotoxicity was assessed using a single target cell line and fixed assay conditions, which may not capture performance across different antigen densities or tumor contexts. Third, mechanistic interpretation remains limited because we did not dissect how individual substitutions alter receptor biophysics or downstream signaling pathways.

Our results also underscore the limitations of relying solely on sequence-based fitness predictors to engineer synthetic receptors. We used EVmutation to prioritize substitutions predicted to be evolutionarily tolerated, which likely reduced the number of catastrophic loss-of-function variants. However, evolutionary fitness does not necessarily translate to CAR-specific properties such as immune synapse formation, receptor clustering, or signaling dynamics in primary T cells. Therefore, computational prioritization should be viewed as a strategy to focus library design rather than a substitute for functional screening.

In summary, we demonstrated that a focused, fitness-guided point-mutant library combined with mRNA electroporation enables scalable, primary T-cell–based screening of CAR variants. This workflow identified candidates with improved cytotoxicity, and subsequent lentiviral validation revealed that #306 may provide an intrinsic functional advantage while preserving a CD62L^high^ phenotype. These findings support the utility of arrayed transient screening as a rapid initial step for CAR optimization.

## Materials and methods

### Sequence design

The EVcouplings (Hopf et al., 2019) was used to generate multiple sequence alignments (MSAs) and train the EVmutation model for fitness predictions. The amino acid sequences of hinge and costimulatory domains of CD28 were separately passed into the EVcouplings webapp, and the MSAs were made against UniRef100 database (Suzek et al., 2015) for bitscore 0.3 and 0.5, respectively. All other EVcouplings parameters were set as follows: Alignment threshold type = Bitscore; Search iterations = 5; Position filter = 70%; Sequence fragment filter = 50%; Removing similar sequences = 90%; Downweighting similar sequences = 80%. Fitness of all single-residue substitutions were calculated using the EVmutation model trained on the MSAs. The resultant single mutant matrices were downloaded from the EVcouplings webapp. The substitutions with higher fitness for each domain were selected to make domain variants. Double-stranded DNAs encoding all possible combinations of the hinge domain variants, transmembrane domain and co-stimulatory domain variants were synthesized as eBlocks Gene Fragments by Integrated DNA Technologies.

### Cell culture

293FT (Thermo Fisher Scientific) was cultured with DMEM (Nacalai tesque) with 10% FBS (Corning), 2 mmol/mL L-glutamine, 100 U/mL penicillin, 100 µg/mL streptomycin, and 1 mmol/mL sodium pyruvate (Nacalai tesque). Nalm6 cell was obtained from RIKEN BRC and stably transduced with EGFP and firefly luciferase to establish Nalm-Luc cell using the lentiviral vector pHR-SFFV-Luc-p2A-EGFP. Nalm-Luc cell was cultured with RPMI1640 (Nacalai tesque) supplemented with 5% FBS, 2 mmol/mL L-glutamine, 100 U/mL penicillin and 100 µg/mL streptomycin. A375 was obtained from ECACC and stably transduced with truncated CD19 using the lentiviral vector pHR-SFFV-tCD19 (A375-CD19). A375-CD19 was cultured with DMEM with 10% FBS, 100 U/mL penicillin, and 100 µg/mL streptomycin.

Primary human CD8 T cells (STEMCELL Technologies, Vancouver, BC, Canada) were cultured in X-VIVO15 medium (LONZA, Basel, Switzerland) with 5% human serum (Sigma-Aldrich, St. Louis, MO, USA), 100 μM 2-mercaptoethanol (FUJIFILM-Wako Pure Chemical Corporation, Osaka, Japan), 25 U/mL human IL-2 (Peprotech, Rocky Hill, NJ, USA), 10 ng/mL human IL-7 (Miltenyi Biotec, Gladbach, Germany), and 10 ng/mL human IL-15 (Miltenyi Biotec).

### Plasmid construction

The dsDNAs were synthesized by IDT and individually subcloned to the BamHI digested backbone (FMC63-(BamHI)-CD3z-P2A-EGFP) with NEBuilder HiFi DNA Assembly Master Mix. We transformed the cloning product into HST08 competent E. coli cells (Takara). Transformed cells were plated LB agar plate with carbenicillin. Plasmid purification and sequencing were performed by AZENTA.

### Preparation of CAR T cells using electroporation

The constructed plasmids were used as templates for PCR to introduce T7 promoter (ggatccTAATACGACTCACTATAgggataat) (Conrad et al., 2020) for in vitro transcription. mRNAs were prepared by BEX using a T7 RNA polymerase-based in vitro transcription reaction incorporating pseudouridine-5’-triphosphate (pseudo-UTP). For co-transcriptional 5’ capping, anti-reverse cap analog was used. Primary human CD8+ T cells were activated using Dynabeads Human T-cell Activator CD3/CD28 (ThermoFisher) at 25 µL per 1×10^6^ T cells. Two days after activation, Dynabeads were removed using a magnet stand (Thermo Fisher Scientific), and then 2×10^5^ CD8+ T cells were electroporated with 500 ng mRNA using 384-well Nucleofector System (Protocol EO-115, Lonza). After electroporation, CD8+ T cells were cultured with X-VIVO15 (Lonza) supplemented with 5% human AB serum (Millipore-Sigma) and 100 µM 2-mercaptoethanol (FUJIFILM Wako Chemicals). Pipetting operation was performed using Biomek i7 (Beckman Coulter).

### Transduction

HEK293FT cells (Thermo Fisher Scientific) were transfected with the pHR-GOI vector and packaging plasmid mix (pMD2.G, pCMVR8.74, and pAdvantage) to produce an incomplete replication lentivirus vector. The viral supernatant was harvested at day2 and concentrated with Linti-X concentrator (Takara). Human CD8+ T cells were stimulated with Human T-activator CD3/CD28 Dynabeads (Thermo Fisher Scientific) at a 1:1 cell-to-bead ratio, cultured for 24 hours, and mixed with the concentrated viral supernatant. After culturing for 2 days, the beads were removed from the T cell culture using a Magnum FLX magnet plate (ALPAQUA,). After 3 days, transduction efficiency was measured by detecting GFP-expressing cell rate using flow cytometer (CytoFLEX S, Beckman Coulter). Transduced cells were used for cytotoxic assays.

### Cytotoxicity assay

For mRNA CAR T cell screening, the cytotoxicity assays were performed using a bioluminescence-based method. Briefly, Nalm-Luc cells were seeded into 4 wells of a 384-well round-bottom plate (Sumitomo Bakelite). Subsequently, CAR T cells were added at an effector-to-target (E:T) ratio of 1:2. After co-culturing for 23 h, the luciferin reagent (BrightGlo, Promega) was added at half the volume of the total culture medium, and luminescence was measured using a microplate reader (COLONA SH-9000).

Target cell cytotoxicity was determined using the following formula:

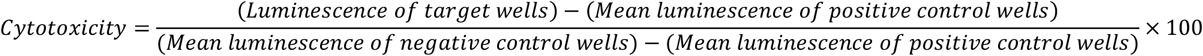

Negative control wells contained only Nalm-Luc cells, while CAR T cells genetically engineered to express wild-type CAR were added to positive control wells at an E:T ratio of 2:1. Luminescence was measured in 16 wells for each control, and the mean luminescence value was used to calculate cytotoxicity. The cytotoxicity values for each variant from four wells were converted to mean values. To validate cytotoxicity of lentiviral transduced CAR T cell, GFP+ CAR T cells were sorted and then cocultured with Nalm-Luc cells for 40 hr at an E:T ratio of 1:16. Luminescence was detected by a microplate reader (COLONA SH-9000).

### CD62L expression analysis

Lentiviral transduced CAR T cells were sorted to enrich CAR+ (GFP+) cells. The sorted CAR T cells were cocultured with CD19-expressing A375 at an E:T ratio of 1:2 for 48 hours. The cocultured cells were stained with PE-conjugated CD8 (BioLegend) and Brilliant Violet 421-conjugated CD62L (BioLegend). Flow cytometry was performed using the CytoFLEX S, and the data were analyzed FlowJo version10 (BD Bioscience).

## Authors contributions

A.O. designed the experiments. S.H. designed the CAR variant sequences. A.O. and Y.I. performed the experiments. A.O., S.H. and Y.I. analyzed the data. S.H. and A.O. created the figures and wrote the manuscript.

## Declaration of competing interest

The authors are currently full-time employees at Hitachi, Ltd.

## Acknowledgments

The authors thank Satoru Mimura for technical assistance, Shinsuke Araki for helpful discussion, Shizu Takeda and Hiroko Hanzawa, for advice, encouragement, and support.

